# A Model-Free Approach for Detecting Genomic Regions of Deep Divergence Using the Distribution of Haplotype Distances

**DOI:** 10.1101/144394

**Authors:** Mats E. Pettersson, Marcin Kierczak, Markus Sällman Almén, Sangeet Lamichhaney, Leif Andersson

## Abstract

Recent advances in comparative genomics have revealed that divergence between populations is not necessarily uniform across all parts of the genome. There are examples of regions with divergent haplotypes that are substantially more different from each other that the genomic average.

Typically, these regions are of interest, as their persistence over long periods of time may reflect balancing selection. However, they are hard to detect unless the divergent sub-populations are known prior to analysis.

Here, we introduce HaploDistScan, an R-package implementing model-free detection of deep-divergence genomic regions based on the distribution of pair-wise haplotype distances, and show that it can detect such regions without use of *a priori* information about population sub-division. We apply the method to real-world data sets, from ruff and Darwin’s finches, and show that we are able to recover known instances of balancing selection – originally identified in studies reliant on detailed phenotyping – using only genotype data. Furthermore, in addition to replicating previously known divergent haplotypes as a proof-of-concept, we identify novel regions of interest in the Darwin’s finch genome and propose a plausible, data-driven evolutionary history for each novel locus individually.

In conclusion, HaploDistScan requires neither phenotypic nor demographic input data, thus filling a gap in the existing set of methods for genome scanning, and provides a useful tool for identification of regions under balancing selection or similar evolutionary processes.

## Background

In comparative genomics, there has, in addition to the search for selective sweeps, long been an interest in detecting genomic regions that have a deeper divergence time than the majority of the genome, as such regions are likely to reflect interesting evolutionary processes. In particular, balancing selection and introgression events have attracted considerable interest, but have proven difficult to detect, as they lack the distinctive footprint - loss of heterozygosity across the population - of a hard selective sweep. This property also impedes analyses that rely on comparing sub-populations, such as F_ST_, as a locus under balancing selection is expected to segregate within each subpopulation. Similar problems affect coalescent methods, such as the one proposed by DeGiorgio et al. (DeGiorgio, et al. 2014), which rely on recombination data as well as on explicit assumptions about the demographic model underlying the regions of interest, which also have limited scalability due to their complexity. All in all, balancing selection is still challenging to detect and all current methods have some associated issues. For a recent review of the state of the field, see Fijarczyk and Babik (Fijarczyk and Babik 2015).

However, the ability to sequence many individuals at a reasonable cost has led to progress in this field, and to date there are several examples in the literature that use differences between populations or related species to identify divergent regions, typically linked to population specific adaptations e.g. (Huang, et al. 2012; Rubin, et al. 2012; Guo, et al. 2013; Varshney, et al. 2013). A common theme is that the analyses rely on tests, such as F_ST_ or χ^2^ on allele frequencies, between pre-defined population groups. This is a powerful approach, but it is not suited for loci that segregate within populations, as in the case of balancing selection, or loci that have been selected according to unknown population structure. Traditionally, measures like average heterozygosity or Tajima’s D (Tajima 1989) have been used to detect signals of selection in sample sets lacking these known contrasts, but these metrics are aimed at signals that are consistent across the majority of samples, e.g. selective sweeps.

In addition to the above mentioned contrast based measures, there are methods relying on local phylogenies ((Zamani, et al. 2013); (Geneva, et al. 2015)) that allow for detection of regions with evolutionary histories that differ from the genome in general. These share some features with looking for deep divergence. Here, we employ the same basic idea of using local patterns among chromosomes in a sample to propose a simple, computationally efficient method that allows detection of deep-divergence regions in a model-free, un-supervised fashion. Using only phased genotypes as input data, datasets of a hundred or more whole-genomes of approximately 1 Gb size can be analysed in a matter of hours on a desktop computer. Our method relies on the common feature that regions of deep divergence have at least one group of haplotypes which is much more similar within itself than compared to the remaining set of haplotypes in the population. This situation will cause the distribution of all pairwise nucleotide distances to be, at least, bi-modal. This multimodality is due to the short within-group distances being separated from the out-of-group or, if there are two distinct haplotypes, across-group distances. Fig. 1 provides a cartoon representation of the idea behind the method.

**Fig. 1.**
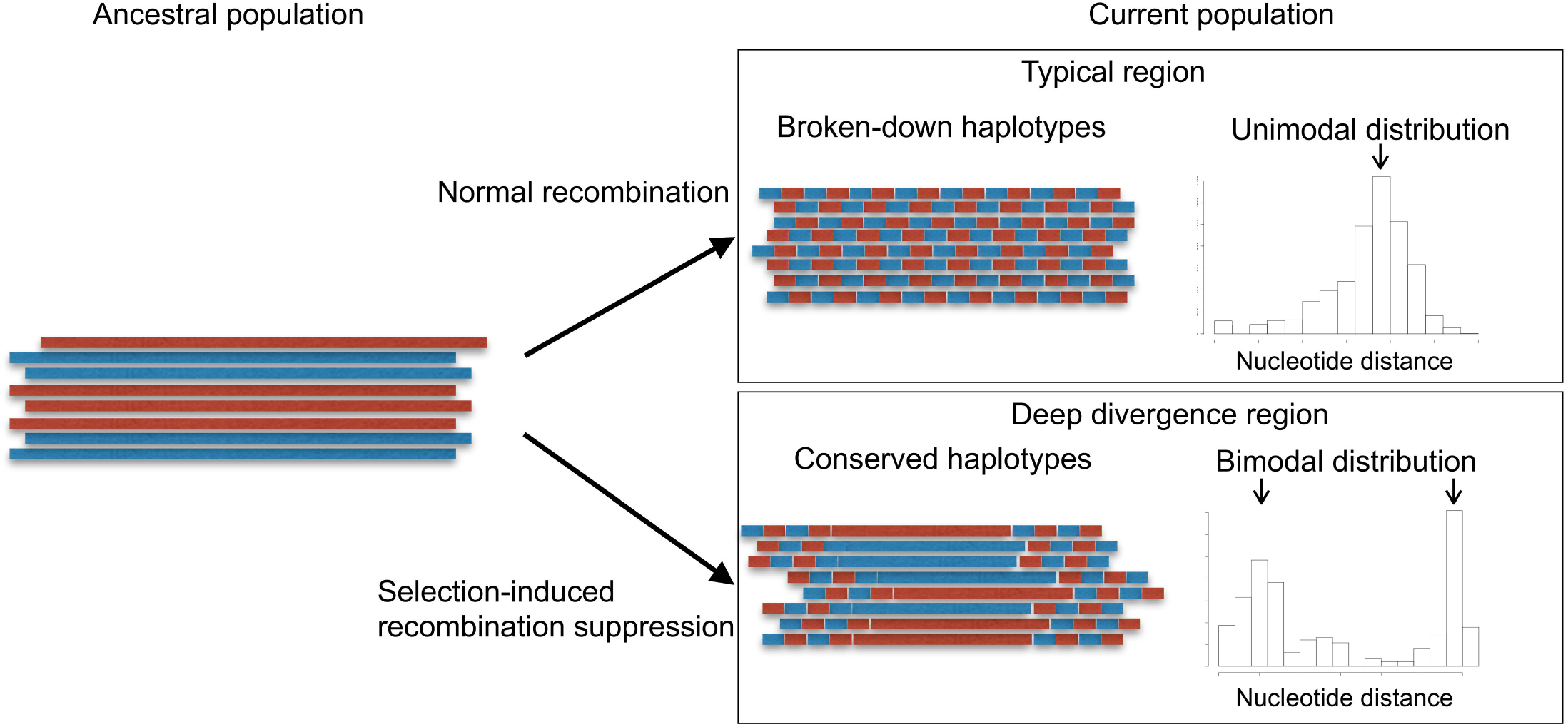
Schematic presentation of the basis for the haplotype distance method. The cartoon shows the progression of two hypothetical genomic regions from an ancestral state. The upper path is a typical region experiencing mixing through recombination, the lower path a region where recombination is suppressed, through selection on long haplotypes, re-arrangements or other means.

From this observation, it follows that statistics that examine the nucleotide distance distribution, in particular its modality, can be used to detected regions of interest. Specifically, Hartigan’s diptest (Hartigan and Hartigan 1985) can be applied, as it measures deviation from unimodality. Also simpler summary statistics, such as the standard deviation or the range of the distribution can be informative.

As outlined in the introduction, the key advantage of the proposed approach, compared to population-contrast based methods, lies in the fact that it does not make any *a priori* assumptions about which samples are similar, although it is a requirement that the frequency of the rare haplotype class is sufficiently high to influence the distribution as a whole. Since each window is examined independently, it is also possible to catch different patterns in different parts of the genome. The lack of need for a specific contrast is especially advantageous in situations where little useful auxiliary information beyond the genotypes is known, which is typically the case when looking for natural selection since our understanding of the components of fitness is very limited for most species.

## Results

### Case study I: Ruff

#### Outline

The ruff (*Philomachus pugnax*) is a Palearctic wader with a spectacular lekking behaviour, as well as an intricate mating system comprising three types of males, *Independents, Satellites* and *Faeders*, with different mating strategies. Using a dataset of 25 sequenced ruff individuals (15 *Independents*, 9 *Satellites* and a single *Faeder*), Lamichhaney et al. (Lamichhaney, Fan, et al. 2016) identified a 4.5 Mb inversion that is associated with the two alternative morphs, *Satellite* and *Faeder*. The haplotypes are remarkably different, with roughly 1.4% nucleotide divergence, indicating that they split about 3.8 million years ago. The inversion is a recessive lethal because it disrupts an essential gene, which means that *Independents* are all homozygous *I/I, Satellites* and *Faeders* are heterozygous *I/S* and *I/F*, respectively. Based on the previous study, it is known that this locus is an archetypical example of a deep divergence region under balancing selection, and thus a good proof-of-concept test for the haplotype distance method. Below we show that, in contrast to other methods using only genetic data, HaploDistScan recovers this signal without utilising either demographic or phenotypic data.

#### Results

We used the previously published data and screened the genome using 100 kb windows, and detected a highly significant signal perfectly overlapping the 4.5 Mb inversion on scaffold 28, without using any *a priori* phenotypic information (Fig. 2*a*). In contrast, Tajima’s D and average expected heterozygosity, standard measures commonly used to detect selection in the absence of *a priori* informative contrasts, do not provide any detectable signal in this region (Figs 2b and c). Fig. 2*d* contains a detailed visualisation (distance histogram, heuristic clustering tree, and heatmap) for one of the windows in the inversion peak, showing that *Independents* and *Satellites* are clearly distinct in this genomic region. We see that *Satellites* and *Faeders* are not perfectly resolved into the expected heterozygote pattern – no haplotypes closely resemble each other in these groups. This is most likely due to technical difficulties in accurately phasing the very long heterozygous stretches with high divergence, leading to mix-up between the chromosomes in each individual. While this issue is external to HaploDistScan, it could affect signal detection. However, as can be seen in Fig. 2*a*, the haplotype distance scan method is robust to these phasing issues, and detects the signal caused by the distinct *Independent* group. The distances within this group are in the order of 400 SNP per 100 kb whereas the between-group distances are about 3-fold higher. Lamichhaney et al. (Lamichhaney, Fan, et al. 2016) showed that the *Satellite* haplotype arose about 500,000 years ago due to a rare recombination event(s) between an *Independent* and *Faeder–like* haplotype and thus appears as a chimeric form of these chromosomes. In the 100 kb window presented here it is most closely related to the *Faeder* haplotype. The haplotype distance scan reveals several other, shorter and less extreme, regions of deep divergence (Fig. 2*a*), that, while currently not linked to any known phenotype, provide information that can aid future genetic studies of ruff populations.

**Fig. 2.**
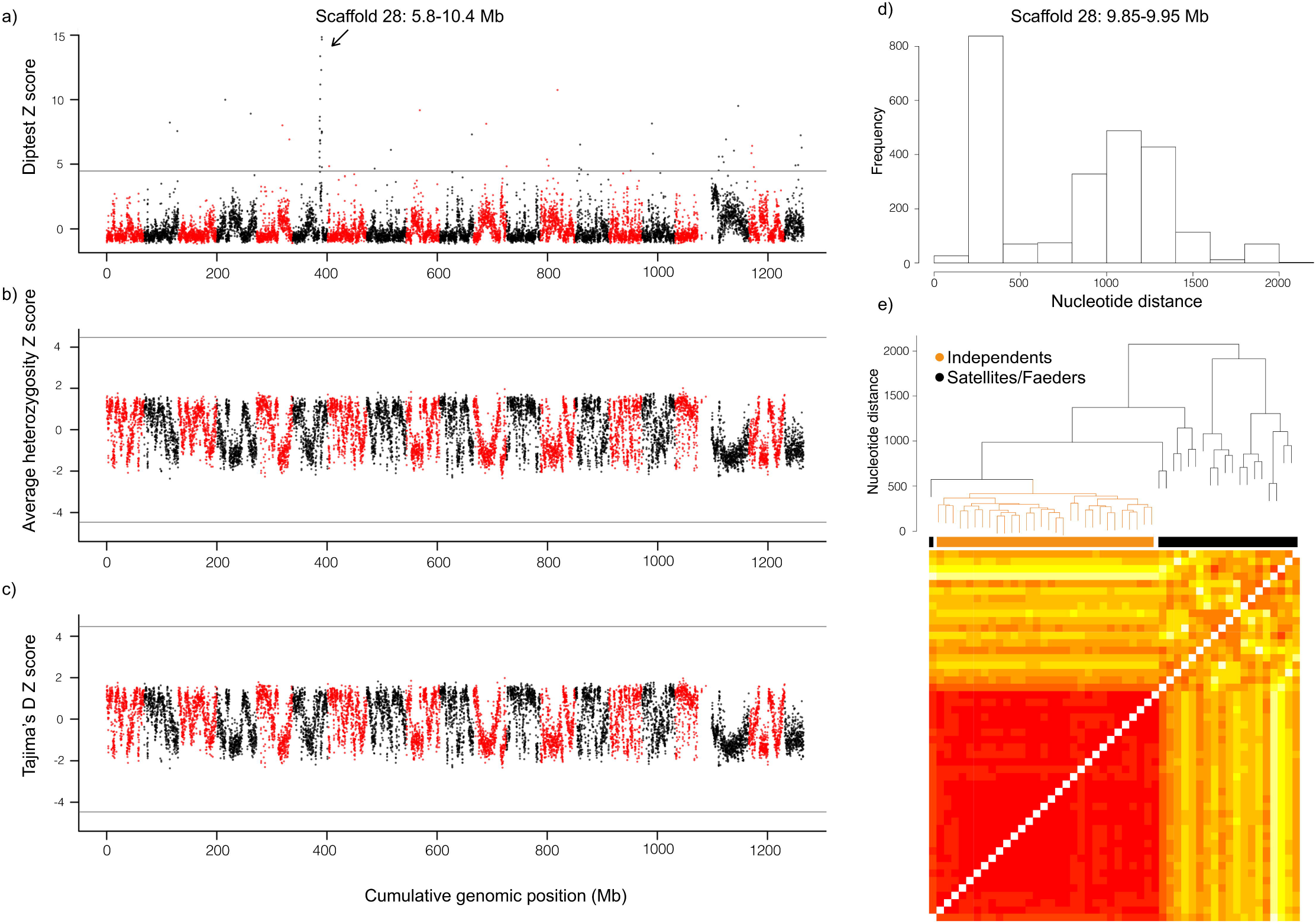
Three genome-wide scans of the ruff genome. Each panel displays one metric, calculated at each 100 kb window across the Ruff genome. All windows containing less than 200 SNP have been filtered out. Panel *a* contains the dip-test statistic calculated on the haplotype distance distribution. Panel *b* is average heterozygosity and panel *c* is Tajima’s D. In order to facilitate comparisons, all metrics has been transformed into Z values, as described in the “significance” section. Panel *d* shows the HaploDistScan output for the top window from the inversion peak, with colour indicating the “*Independent*” and “*Satellite*” clades, respectively.

### Case study II: Darwin’s finches

#### Outline

Recently, Lamichhaney et al. (Lamichhaney, et al. 2015) presented a phylogenetic analysis based on 120 whole-genome sequenced birds representing all the 14 species of Darwin’s finches inhabiting the Galapagos archipelago and Cocos island. Besides providing new insight about the phylogenetic relationships in this recent adaptive radiation of birds, the study revealed a major locus underlying differences in beak shape, located from 320 to 560 kb on scaffold JH739921, and showed that the likely causative gene is *ALX1*, which encodes a transcription factor with an established role for craniofacial development. The two haplotypes associated with blunt and pointed beaks showed a deep evolutionary divergence (estimated at 900,000 years before present), much further back than the genomic average. Thus, this locus constitutes a second previously identified case of a region under balancing selection.

#### Results

We performed a genome scan, with a window size of 100 kb, for a subset of 39 birds representing the different species of ground finches (Fig. 3*a*), and the haplotype distance scan method detected the *ALX1* locus without using any *a priori* phenotypic information. A detailed analysis of the signal recovers the result from (Lamichhaney, et al. 2015), as the sharp- and blunt-beaked birds fall in separate halves of the haplotype tree (Fig. 3*b*).

**Fig. 3.**
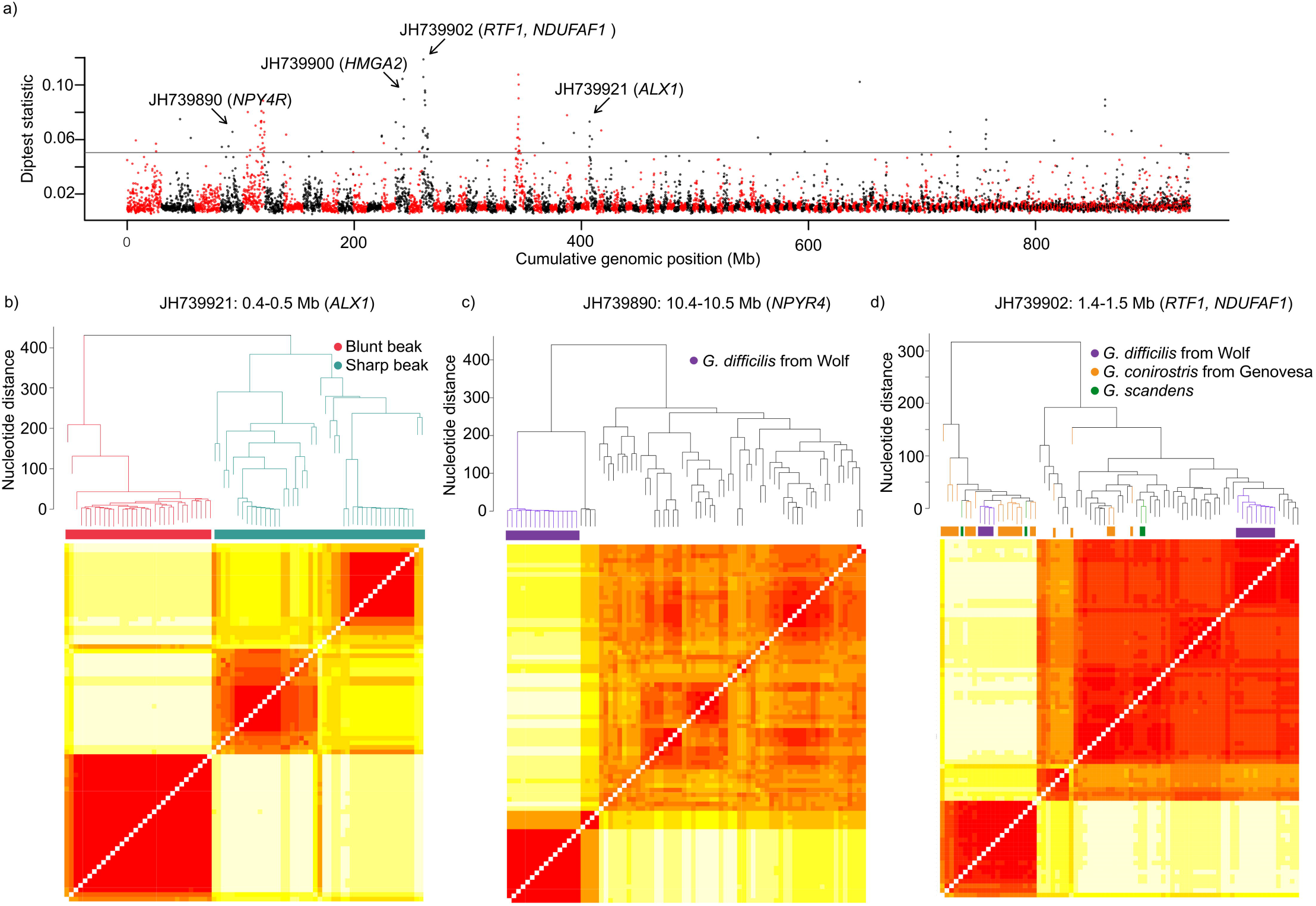
Significant regions in the Darwin’s finch genome. Panel *a* shows the diptest statistic scan across the entire *G. fortis* reference genome. The window size is 100 kb, and all windows containing less than 200 SNP have been filtered out. The colour alternates with each scaffold, and the x-axis is the cumulative physical position. Panels *b* through *d* each displays one significant 100 kb region. In the heat maps, red indicates high similarity (short edit distance), while pale yellow means low similarity (long edit distance). The regions are JH739921:0.4-0.5 Mb (*b*), JH739890:10.4-10.5 Mb (*c*), and JH739902:1.4-1.5 Mb (*d*).

In the same data set, we were also able to identify a deep haplotype divergence involving the *Geospiza difficilis* population on Wolf island at position 10.5 Mb on scaffold JH739890, where the birds from the Wolf island are fixed for a haplotype that has no close relative in the rest of the data set (Fig. 3*c*). Extending the analysis to the entire data set reported previously (Lamichhaney, et al. 2015) (Supplementary Fig. S1) reveals that this haplotype is shared with *G. difficilis* birds from the nearby Darwin island (clade 1a), while the *G. difficilis* samples from Pinta, Santiago and Fernandina form the next-nearest group (clade 1b), together with one *G. scandens* bird and one *G. fortis* bird. This pattern is not typical for the genome as a whole, where these *difficilis* populations do not form a monophyletic clade, but supports the idea that the Darwin/Wolf *difficilis* populations are of mixed ancestry (Lamichhaney, et al. 2015). Furthermore, the presence of the “*difficilis*” haplotype in one *G. scandens* and one *G. fortis* individual, indicates some amount of gene flow into those populations as well. The haplotype distribution makes the region a promising candidate for any trait that is shared across the different *G. difficilis* populations, excluding the *G. difficilis* from Genovesa.

This region contains the *NPY4R* gene (coding for neuropeptide Y receptor Y4), which suggests a possible association to behaviour, as neuropeptide Y has been linked to several behavioural traits (Tatemoto 2004). Specifically, food intake is a trait previously linked to *NPY4R* that is likely to have a fitness effect in these populations, since *G. difficilis* from Darwin and Wolf share the habit of feeding on blood from sea-birds (Schluter and Grant 1984) and are thus commonly referred to as “vampire finches.”

A distinguishing feature of the HaploDistScan method is the ability to detect regions of the genome where divergent haplotypes are segregating within several species, as *a priori* divisions of the data set cannot disrupt it. An example of such a region is found on scaffold JH739902 (Fig. 3*a*) where a segregating haplotype is found at intermediate frequencies in three different ground finches (*G. conirostris, G. difficilis* and *G. scandens;* Fig. 3*d*). This region contains the *RTF1* (Paf1/RNA polymerase II complex component homolog) gene, affecting transcript elongation, as well as the *NDUFAF1* (NADH dehydrogenase (ubiquinone) complex I, assembly factor 1) gene, a component of mitochondrial complex I.

Supplementary Fig. S2 shows the same region, but includes the full dataset from Lamichhaney et al. (Lamichhaney, et al. 2015). In this extended data set, there is clear support for one of the haplotypes segregating within several ground finch species sharing ancestry with the version found in present day tree finches; a clear example of where the local sequence tree does not match the species tree based on the entire genome. This indicates an introgression event occurring not too long before the radiation of the current tree finch species, since the two versions of the haplotype form separate sub-clades (clades 2c and 2d, respectively). The presence of two *G. conirostris* haplotypes inside the tree finch sub-clade (2d) could indicate a second, more recent, introgression. A second event that persists over time further supports the idea that this haplotype confers a fitness benefit in the recipient species, at least under some conditions.

The other segregating haplotype (clade 4), which is fixed in the other ground finch populations, is also found in the otherwise more distant Cocos finches (clade 4a). This pattern suggests lateral transfer of genetic material, since Cocos finches are, in the general phylogeny, an outgroup to tree- and ground finches. However, a similar pattern could occur if instead all other finches (i.e. clades 1-3) were carrying very old haplotypes. In order to distinguish the two scenarios, we calculated the between-species sequence divergence (d_XY_) for this locus and compared with the genome in general, with results shown in Supplementary Fig S3. Given that the Cocos finch and ground finches are more similar in this region than the genomic average, while the region is typical when compared to an outgroup species, an introgression event is the more likely explanation. As with the clade 2-haplotype, the Cocos finches and ground finches form separate sub-clades, placing the transfer event at a time point before the radiation of the present day species.

In addition, the full data set adds three new haplotype groups to the pattern. One, distantly related to the tree finch haplotype, is found in *G. difficilis* from Pinta, Santiago and Fernandina (clade 2a) together with *Platyspiza crassirostris* (clade 2b). Notably *G. difficilis* from Genovesa do not carry this haplotype, but are instead found among the remaining ground finches (clade 4), an observation consistent with previous results (Lamichhaney, et al. 2015), where this population rarely grouped with the other *G. difficilis* birds; *G. difficilis* from Genovesa were in fact proposed to be a distinct species and named *G. acutirostris*. The last two haplotype groups are found in the Warbler finches (clades 1 and 3), where the one exclusive to *Certhidea olivacea* (clade 1) is the one furthest removed from all other haplotype groups. As can be seen in Fig. 3*d*, the nucleotide distances between the different haplotypes in this locus are similar to those found at the *ALX1* locus, indicating a similar age. Thus, the fact that they are still segregating in several observed species is a strong indication of balancing selection.

Beyond the examples discussed above, several additional deep-divergence regions were identified (Fig. 3*a*). For example, the region around the gene coding for High-mobility group AT-hook 2 (*HMGA2*) on JH739900 (6.0 - 7.4 Mb), was recently shown to influence beak size (Lamichhaney, Han, et al. 2016). The full list of significant regions, several of which may warrant further investigation, can be found in S1 Table.

### Simulation-based validation

In order to estimate power of detection as a function of divergence time of a deep-divergence region, we performed a simulation study using data generated by the coalescent simulation software “ms” (Hudson, 1992). We simulated a genome consisting of 1000 independent regions, 20 of which were designated as deep-divergence loci. The divergence time (*t*; in fractions of 4Ne) for these inserted target regions was evenly spaced from 0 to 1. In addition, in order to estimate the effect of background stratification, we varied the recombination frequency across each locus (the “*r*” parameter in “ms”) from 100 to 10, where lower values indicate less recombination and thus more stratification in the sample set. However, even the value of 100, which corresponds to an effective 0.1 cM/Mb under the condition used, is lower than observed in most organisms, and can thus be considered to represent a non-ideal situation where recombination is impaired by population structure. The simulation was repeated 100 times for each set of parameters, allowing us to calculate the capture rate, in terms of the dip-test statistic exceeding the threshold, as well as the number of false positives (S5 Figure). As can be seen in panel *a*, the power is high in the upper right corner of the plot, with the conditions *t* >= 0.5 and *r* >= 40 roughly bounding the area with close to 100 % power. Panel *b* shows that the false negative rate peaks at 0.4% (4.2 hits out of 980 possible) for *r* = 20, which indicates that the background-based threshold calculation efficiently controls the absolute number of false positives in stratified conditions. The number drops to close to zero for *r* >= 50, which, combined with the high power in this region, gives a low ratio of false-to-true positives (panel *c*).

## Discussion

As shown in the case studies above, there are signals of selection and divergence that are difficult to detect in the absence of information beyond individual genotypes. In particular, this is true for signals that are not consistent across whole populations, such as balancing selection. Consequently, auxiliary phenotypic and/or demographic information has been needed to detect such loci.

However, in the cases studies described above, we show that the haplotype distance method provides a robust means to detect such signals in an un-supervised manner using only genotypic information. We were able to identify known signals in both ruff and Darwin’s finches, while in addition also detecting several novel regions, presumably related to hitherto unexplored phenotypes, which warrant further examination.

The Darwin’s finch study in particular highlights the usefulness of the method in a complex sample, where recurring transfer of genetic material between otherwise isolated populations has resulted in different patterns of relationship in different genomic regions. HaploDistScan allows us to treat these regions independently, and detect potentially conflicting signals in a single run. In addition, the visualisation tools provided in the package make it possible to dissect each signal, and suggest possible explanations based on the contrasts that emerge from the data.

## Conclusions

In conclusion, the haplotype distance method can handle sizeable datasets of whole-genome sequenced individuals, can be applied at an early stage since only phased genotypic data is required, and has the power to detect signals of deep divergence that are not recognizable by other model-free approaches.

## Methods

### Summary of the haplotype distance method

The genome to be analysed is split into consecutive, non-overlapping windows, which are scored independently. The optimal window size to give a reasonable power to detect deviations from unimodality will depend on the haplotype size in the population as well as SNP density. In general, it is the case that shorter windows lead to increased inter-window variance and thus to reduction in power. The benefit is that, naturally, smaller window size gives higher resolution and thus possibility of detecting shorter signals. For each window, all haplotypes are extracted from the genotype matrix, and their pairwise edit distances, i.e. the number of nucleotide changes, are calculated. The distribution of this set of distances determines the score for the corresponding window.

### Metrics

We propose three possible metrics to characterise the distance distributions, each with their own pros and cons:

1. Hartigan’s diptest statistic (Hartigan and Hartigan 1985), which measures deviation from unimodality. We use the statistic rather than the P-value, as we want to estimate significance from the genomic distribution rather than against a null hypothesis of no deviation from uni-modality, in order to compensate for the level of structure present in the analysed data set. This metric works well for cases with one or two distinct haplotypes, but has lower discriminatory power in situations where three or more groups cause distributions to overlap and, thus, smoothen out the overall distribution.
2. The standard deviation of the distance distribution divided by the total number of SNPs in each region. This measure is, on average, less powerful than the diptest statistic, as factors that introduce skew to the distribution will also increase the standard deviation. However, it works better in windows with several distinct haplotype groups, and thus complements the above test.
3. The range (difference between the largest and the smallest distance) divided by the total number of SNPs in each region. This metric has the lowest average power, as it does not include any mechanism to suppress noise, but has the advantage of being frequency-independent. It is therefore most useful for identifying haplotypes that are very different from the others in the population, but sufficiently rare to be present in only one or two copies in the sampled data set.

### Significance testing

Given that the method aims at finding regions that are distinct from the genome in general, it makes sense to make use of the genomic distribution in order to determine significance. This has the benefit of accounting for both structure in the dataset, as well as the average level of divergence.

We apply a procedure similar to a z-transformation by calculating the mean and standard deviation of the distribution across all windows for each metric, and put the threshold where a normal distribution with this parameters would have its quantile corresponding to 1-0.05/*N*, where *N* is the number of windows. Thus, if the distribution across windows is normal we have a 5% risk of detecting a single false positive result in each genome scan.

While this procedure provides correction for the genomic level of variation, the consequence is that power is decreased in datasets with strong structure. Therefore, it is typically best to use a dataset with as little population structure as possible, and analyse known population strata separately.

### Software

In order to facilitate the analysis described above, we have implemented the procedure in an R (R Core Team 2015) package: HaploDistScan which provides a set of functions needed for performing the procedure described above. The package reads in and processes a phased Variant Call Format (VCF) file, which might be generated from GATK (McKenna, et al. 2010) and BEAGLE (Browning and Browning 2013), and allows the entire analysis to be completed in a few commands. A simple example is shown in the package vignette, while detailed usage descriptions are found in the package documentation.

### Performance

The R package was able to perform the scan of the ruff data (described above), which covers 25 whole-genome sequenced individuals each genotyped at 16.6 million SNPs, split into 12,600 windows, in approximately one hour on a MacBookPro laptop computer (16 GB DDR3 RAM, 2.8 GHz Intel i7 CPU).

### Empirical method evaluation

#### Window size

The effect of varying the window size in the ruff data set from case study I is shown in Supplementary Fig. S4. In qualitative terms, the major difference is in the background level, which decreases with increasing window size, and that there are a number of single windows that display high divergence in the smaller window size scans. Those signals could be of interest, but, since they are typically supported by fewer SNPs and are not part of a coherent context, some caution is prudent. In general, the pattern of one dominant signal is consistent across both the different window sizes and divergence metrics.

#### Divergence metrics

In the ruff data, the range and standard deviation metrics are highly correlated (*r* = 0.89), whereas the diptest statistic is more distinct (*r* = 0.25 and *r* = 0.30 with range and standard deviation, respectively). This result is consistent with the reasoning in the metrics section indicating that the metrics are tuned for different versions of deep divergence signals, and is thus likely to be the typical metrics’ behaviour. Consequently, it can be recommended to examine at least the diptest statistic and standard deviation metrics in order to find most regions of interest.

### Data deposition

HaploDistScan is available for download at: https://github.com/HaploDistScan/HaploDistScan It is an R package, distributed under the GPL license. The reads underlying the analyzed data sets are available at NCBI. They can be found at http://www.ncbi.nlm.nih.gov/sra?term=PRJNA263122 (Darwin’s finches) and http://www.ncbi.nlm.nih.gov/sra?term=SRA266458 (ruff). Genotype calls (VCF files) for all individuals are available upon request.

## Funding

This works was supported by the European Research Council program Bateson (to LA). MK was supported by the European Research Council and by the Swedish Foundation for Strategic Research. The funding agencies had no involvement in the design of the study, the implementation of the software, analysis of data or writing of the manuscript.

## Authors’ contributions

Designed the study: LA, MP. Implemented the software: MP, MSA, MK. Tested the software: MK, MSA. Performed bioinformatics analysis: MP, SL. Wrote the paper: MP, LA, MSA, MK, SL.

## Acknowledgements

We thank Peter and Rosemary Grant for constructive discussions of the biological interpretation of the Darwin’s finch case study results.

## Supporting Information

**Fig S1. Extended analysis of the JH739890:10.4-10.5 Mb region in the Darwin’s finch genome**. The haplotype visualisation includes the entire dataset from Lamichhaney et al. (Lamichhaney, et al. 2015) for the region displayed in Fig. 3*c*. The distinct clades have been enumerated, and the numbers in the legend indicate in which clades the corresponding population is represented.

**Fig. S2 Extended analysis of the JH739902:1.4-1.5 Mb region in the Darwin’s finch genome**. The haplotype visualization includes the entire dataset from Lamichhaney et al. (Lamichhaney, et al. 2015) for the region displayed in Fig. 3*d*. The distinct clades have been enumerated, and the numbers in the legend indicate in which clades the corresponding population is represented.

**Fig. S3 d_XY_ analysis of the JH739902:1.4-1.5 Mb region in the Darwin’s finch genome**. Panel *a* shows dxy between *Pinaroloxias inornata* (Cocos finch) and *Geospiza magnirostris*. Panels *b* and *c* shows *G. magnirostris* and *P. inornata*, respectively, versus an outgroup species, the tanager *Loxigilla noctis*. Panel *d* shows d_XY_ in 50 kb windows along JH739902, where the black-bordered box indicates the 1.4 - 1.5 Mb window.

**Fig. S4 Examination of the effect of varying window size in the ruff data set**. The figure contains the whole-genome scan across the ruff genome for four different window sizes. 10 kb (a), 50 kb (b), 100 kb (c; default and used in the main text), 500 kb (d).

**Fig. S5 Simulation based validation results**. Panel *a* shows the average fraction of successfully identifying deep-divergence loci across a surface of the divergence time of said locus and the intra-locus recombination rate. Panel *b* shows the number o false positive hits (out of the 980 possible background loci) as a function of the intra-locus recombination rate, while panel *c* shows the false discovery rate, calculated as the number of false hits divided by the number of true hits, also as a function of the intra-locus recombination rate. All values shown are the averages of 100 simulatio runs.

**Table. S1 List of all significant regions in the Darwin’s finch genome**. The table contains the locations and extent of all significantly divergent regions from the Haplotype distance scan of the Darwin’s finch genome. Significant windows less than 500 kb apart have been merged into continuous regions.

